# Direct RNA-seq provides evidence for an antiterminator function of the yeast Hrp1 protein on RNA polymerase II transcripts

**DOI:** 10.1101/2025.09.18.677160

**Authors:** Emma C. Goguen, Kylie A. Zawisza, David A. Brow

## Abstract

Hrp1/Nab4 is an essential nuclear RNA-binding protein that was first identified in the yeast *Saccharomyces cerevisiae* as a cleavage and polyadenylation factor for mRNAs, CF1B, but was later shown to promote termination of some short, noncoding transcripts via the “NNS” termination pathway. Hrp1 binds (UA)_n_ repeats found in both mRNA 3’-UTRs and short noncoding RNA terminators, but its function in 3’-end formation is not fully understood. Our past microarray transcriptome analysis of the heat-sensitive *hrp1-7* allele suggested Hrp1 functions in antitermination of RNA polymerase II (RNAP II) on protein-coding genes. The *hrp1-7* allele has four substitutions and one, M191T, was shown to be primarily responsible for NNS terminator readthrough. Here we show that Hrp1-7 protein has a four-fold and Hrp1-M191T a two-fold decreased affinity for (UA)_5_ in vitro. We used nanopore direct RNA sequencing to assess the transcriptome-wide effects of *hrp1-7* and *hrp1-M191T*, which identified new examples of Hrp1-dependent small nucleolar RNA terminators and mRNA attenuators (regulatory terminators in the 5’-UTR and ORF). For some genes, including *HRP1*, attenuated transcript reads outnumber the corresponding mRNA reads in exponentially growing wild-type cells. We also observed widespread changes in mRNA polyA site selection. In the *hrp1-M191T* strain the polyA site shifts are mostly downstream, while in the *hrp1-7* strain upstream and downstream shifts are more similar in frequency. Our results are consistent with a model in which some of the four substitutions in *hrp1-7* weaken the affinity of Hrp1 for the RNAP II elongation complex, while others interfere with its recognition of specific sequences in nascent transcripts that would normally promote its release from the elongation complex.

## INTRODUCTION

RNA polymerase II (RNAP II) transcribes most genes in eukaryotic genomes and produces both mRNAs and non-coding RNAs. Once RNAP II starts transcribing a gene, regulation of its elongation and termination is conducted mainly by RNA-binding proteins that recognize sequences in the nascent transcript. Termination is coupled with 3’-end formation and, in the yeast *Saccharomyces cerevisiae*, occurs via two major pathways: Cleavage and polyadenylation (CPA) produces mRNAs and the Nrd1-Nab3-Sen1 (NNS) pathway produces both stable and unstable short, non-coding RNAs, including small nucleolar (sno) and small nuclear (sn) RNAs. In the CPA pathway, cleavage and polyadenylation of the nascent transcript is coupled to termination of RNAP II further downstream (Boreikaitė and Passmore, 2023). In the NNS pathway, termination rather than cleavage forms the 3’-end of the transcript, which is polyadenylated by the TRAMP complex and 3’-trimmed or completely degraded by the nuclear RNA exosome (Arndt and Reines 2015).

There are two main models for the mechanism by which RNAP II termination occurs, the allosteric and torpedo models, which are not mutually exclusive (Rodríguez-Molina et al. 2023). In the torpedo model, after cleavage of the nascent RNA the new 5’ end of the polymerase-associated transcript is recognized by the exonuclease Rat1/XRN2, which chases down RNAP II and elicits termination (Connelly and Manley 1988; Proudfoot 1989). There is no evidence for cleavage in NNS termination, but the RNA/DNA helicase Sen1 has been proposed to act in a torpedo-like mechanism whereby it translocates 5’ to 3’ along the nascent RNA to promote RNAP II termination in a manner similar to the bacterial Rho helicase (Steinmetz and Brow 1996; Porrua and Libri 2013; Martin-Tumasz and Brow 2015; Han et al. 2017; Rengachari et al. 2025). The allosteric model proposes that one or more antitermination factors join RNAP II at or near the transcription start site and their subsequent binding to termination signals in the transcript causes their release, reducing the processivity of RNAP II and promoting termination (Logan et al. 1987). Only recently have candidate antitermination factors been identified in mammals, SCAF4 and SCAF8 (Gregersen et al. 2019), which are apparent homologs of the *S. cerevisiae* Nrd1 protein and the *Schizosaccharomyces pombe* Seb1 protein (Yuryev et al. 1996; Steinmetz and Brow 1996; Mitsuzawa et al. 2003). However, an antiterminator function of Nrd1 or Seb1 has not been reported. The yeast protein Npl3/Nab1 has been proposed to have antitermination activity by competing for binding of CPA factors (Dermody et al. 2008).

Hrp1 (also known as Nab4 and CF1B) was initially characterized as a component of the cleavage factor 1 (CF1) complex that binds elements upstream of the polyA site and positions the cleavage and polyadenylation factor (CPF) complex (Kessler et al. 1997, Boreikaitė and Passmore, 2023). Hrp1 recognizes the polyadenylation efficiency element that has a consensus sequence UAUAUA (Guo and Sherman 1996, Kessler et al. 1997, Chen and Hyman 1998). *In vitro* reconstitution of CPA reactions show that Hrp1 is important for cleavage site specificity and in its absence, cleavage occurs at multiple sites (Minvielle-Sebastia et al. 1998, Hill et al. 2019). However, the role of Hrp1 in CPA *in vivo* is still poorly understood.

We previously showed that a heat-sensitive *HRP1* allele, *hrp1-7,* causes readthrough of several NNS terminators including the snoRNA *SNR82* and the attenuator in the 5’-UTR of the *HRP1* gene (Kuehner and Brow 2008, Chen et al. 2017, Goguen and Brow 2023). The *hrp1-7* allele contains four substitutions (Figure 1A), but the strongest effect on NNS terminator readthrough is due to the M191T substitution in the first of two RNA recognition motifs (RRMs), which is predicted to disrupt a hydrophobic cluster that interacts with three bases in its RNA target (Pérez-Cañadillas 2006, Goguen and Brow, 2023). Our previous microarray transcriptome analysis of yeast cells containing the *hrp1-7* allele showed a decrease in full-length mRNA for many protein-coding genes as well as readthrough of several NNS terminators after a shift to restrictive temperature (Chen et al. 2017). These data and a subsequent mutational analysis of *HRP1* led us to hypothesize that Hrp1 acts as an antitermination factor for yeast RNAP II (Goguen and Brow, 2023). While the *hrp1-7* mutation has been shown to have effects on NNS terminators, its effects on polyA site selection in vivo have not been thoroughly investigated.

**Figure 1.**
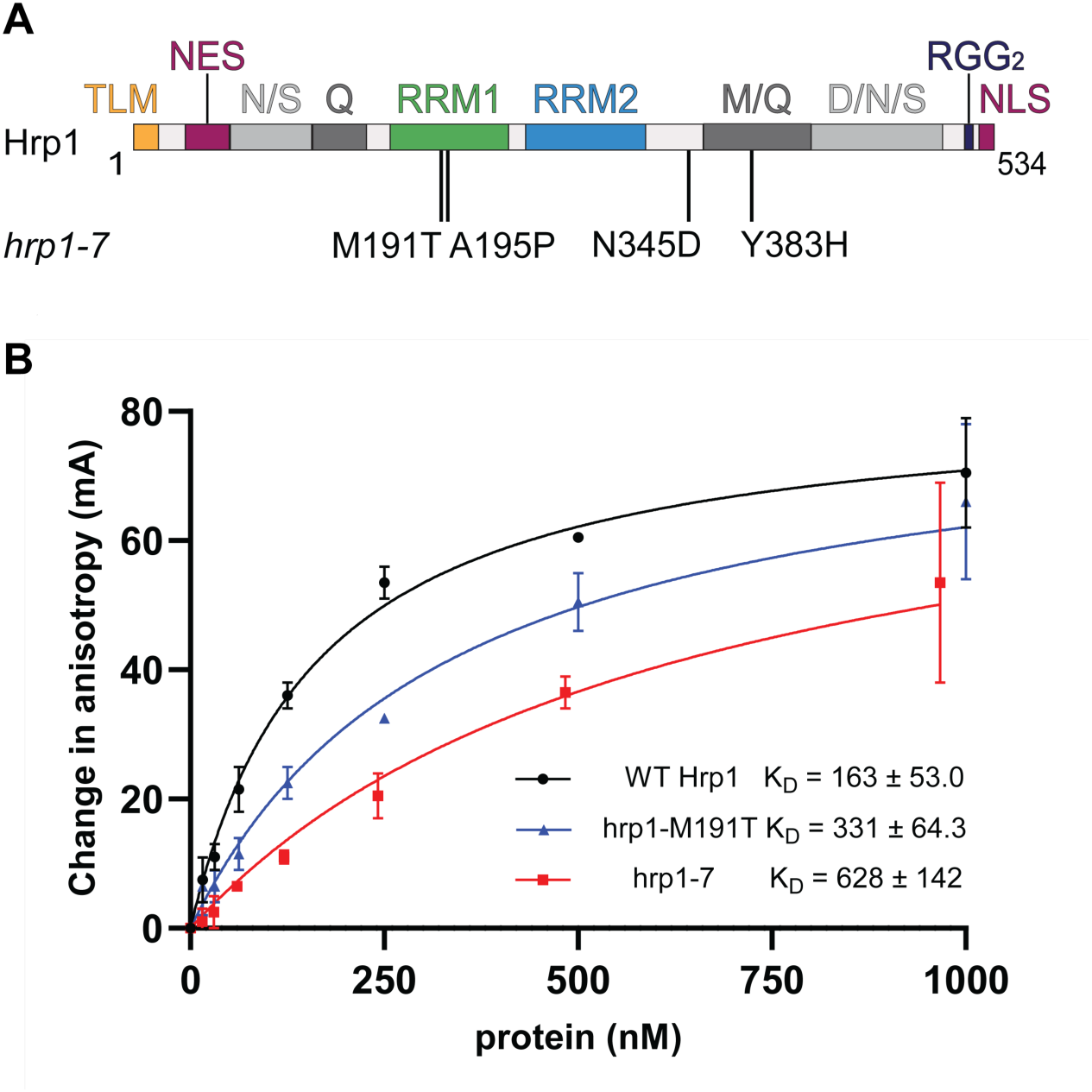
The *hrp1-7* and *-M191T* substitutions decrease affinity for (UA)4 RNA. **A)** Hrp1 contains two central RRMs (green and blue) flanked by regions of low sequence complexity (light and dark gray), named according to their most abundant amino acids (Goguen and Brow, 2023). The *hrp1-7* allele contains the four substitutions shown. **B)** Fluorescence anisotropy of Hrp1 (black), hrp1-M191T (blue), and hrp1-7 (red) protein binding to a FAM-(UA)_4_ RNA probe. Increasing concentrations of protein were incubated with 20 nM of RNA. Each point represents the average ± SEM for two independent experiments. The data points with no error bars had an anisotropy difference of 1 mA. The K_D_ for wild-type and each mutant protein was estimated by fitting data to a one-site binding equation. The error for the K_D_ estimate represents the asymptotic 95% confidence interval of the fit.

To determine the global effects of the *hrp1-7* and *-M191T* mutations on 3’ end formation of mRNAs and non-coding RNAs, we conducted Oxford Nanopore direct RNA sequencing with the wild-type and mutant *HRP1* strains. Consistent with previous data (Goguen and Brow 2023), Hrp1-7 has a stronger effect than Hrp1-M191T on readthrough of the *SNR82* snoRNA terminator, as is also true for seven other snoRNA genes and for the NNS attenuator in the 5’ UTR of *HRP1*. Surprisingly, reads from the attenuated *HRP1* transcripts are more abundant than *HRP1* mRNA reads in all three strains. In contrast, reads from the *NRD1* and *IMD2* attenuated transcripts are less abundant than their corresponding mRNA reads. Using the signature of 3’-truncated mRNA accumulation we identify several other potentially attenuated mRNAs, including the *PAP2/TRF4* mRNA, which encodes the polyA polymerase that adenylates NNS-terminated transcripts. Since Trf4 has been implicated in termination as well as processing, it may autoregulate expression of its mRNA. The broadest effect of the substitutions in Hrp1 is an altered distribution of polyA site use for many protein-coding genes. In the *hrp1-7* strain both upstream and downstream shifts in polyA site selection are common, whereas *hrp1-M191T* more often results in downstream shifts. We propose that, of the four substitutions in the *hrp1-7* allele, some weaken efficiency element recognition, causing downstream shifts in polyA sites, and others weaken interaction with the RNAP II elongation complex, causing upstream shifts.

## RESULTS

### The M191T substitution partially confers the decreased affinity of Hrp1-7 for (UA)_4_

The *hrp1-7* allele contains four substitutions (Figure 1A; Kuehner and Brow 2019). Only the M191T substitution in the first of two RRMs exhibits measurable (albeit weaker) readthrough of the *SNR82* snoRNA terminator on its own (Goguen and Brow, 2023). The M191T substitution is expected to disrupt a hydrophobic cluster that interacts with three bases in its 5’-UAUAUAUA-3’ RNA target sequence, U5, A6, and U7 (Pérez-Cañadillas 2006). To test if Hrp1-7 and Hrp1-M191T have altered affinity for (UA)_4_ RNA, we used fluorescence anisotropy to measure binding to FAM-labeled (UA)_4_ at equilibrium (Figure 1B). Wild-type Hrp1 bound to (UA)_4_ RNA with an apparent dissociation constant (K_D_) of 183 nM, assuming 100% of the protein is active for binding. Hrp1-7 bound with about a 4-fold lower affinity than the wild-type protein, while Hrp1-M191T bound with an affinity about 2-fold weaker than wild-type. Thus, the 2.5-fold stronger readthrough of the endogenous *SNR82* terminator in the presence of *hrp1-7* than *hrp1-M191T* (Goguen and Brow, 2023) is likely due largely to decreased RNA affinity due to one or more of the other three substitutions.

### Direct RNA sequencing detects readthrough of several snoRNA terminators in the *hrp1-7* and -M191T strains

To investigate the global effects of the *hrp1-7* and -*M191T* alleles on RNAP II 3’-end formation, we conducted Oxford Nanopore direct RNA sequencing (dRNA-seq) on total RNA collected from two biological replicates of wildtype *HRP1* and *hrp1-7* cells and one sample of hrp1-M191T cells after shifting from 30 °C to the restrictive temperature 37 °C for one hour. The library preparation method we used ligates sequencing adapters specifically to RNAs with an oligo- or polyadenylated 3’-terminus and thus detects both mature mRNAs and NNS-terminated primary transcripts that have been polyadenylated by the TRAMP complex. Nanopore dRNA-seq starts at the 3’-ends of RNAs, which are therefore mapped with high accuracy, but reads exhibit varying degrees of heterogeneity at the 5’ end, possibly due to RNA degradation (*in vivo* or during extraction) and/or arrested translocation through the nanopore. Nevertheless, transcription starts sites can be deduced for most genes.

For each sample, 1 μg of total cellular RNA was used for library preparation and run on the flow cell for approximately 24 hours, until at least 2 million reads (∼2 billion bases) were sequenced (Supplemental Table S1). The reads were converted to reads per million (RPM) to normalize the samples. The *HRP1* biological replicates have a Spearman correlation coefficient of 0.987 and the *hrp1-7* replicates 0.985 (see Supplemental Table S2 and Supplemental Figure S1 for all pairwise comparisons) indicating that the distributions of reads and genomic coverage are very similar between the replicates. Global analyses were conducted with averaged biological replicates for *HRP1* and *hrp1-7*, while figures show only one replicate for *HRP1*, *hrp1-7*, and *-M191T* that have similar total numbers of mapped reads (2.40 to 2.61 million).

We first examined the dRNA-seq data at loci that previously exhibited *hrp1-7*-dependent readthrough of NNS terminators. Wild-type cells have three different classes of RNAs at and/or just downstream of the *SNR82* snoRNA locus (Figure 2): precursor snoRNAs that are polyadenylated before being 3’ trimmed by the exosome (van Hoof et al. 2000; LaCava et al. 2005), snoRNA terminator readthrough products that span both *SNR82* and the downstream protein coding gene, *USE1*, and *USE1* mRNA. All three RNAs are present in *HRP1*, *hrp1-7, and -M191T* strains, but there is a ∼9-fold increase in the number of *SNR82-USE1* readthrough transcripts in the *hrp1-7* strain relative to wild-type, consistent with our previous studies showing that *hrp1-7* causes strong readthrough of the *SNR82* terminator (Chen et al. 2017; Goguen and Brow, 2023). M191T causes an only ∼3-fold increase in readthrough transcripts, also consistent with our previous findings. There appears to be a distal shift in *USE1* mRNA polyA sites used in *hrp1-7* and *-M191T* compared to *HRP1* (see below).

**Figure 2.**
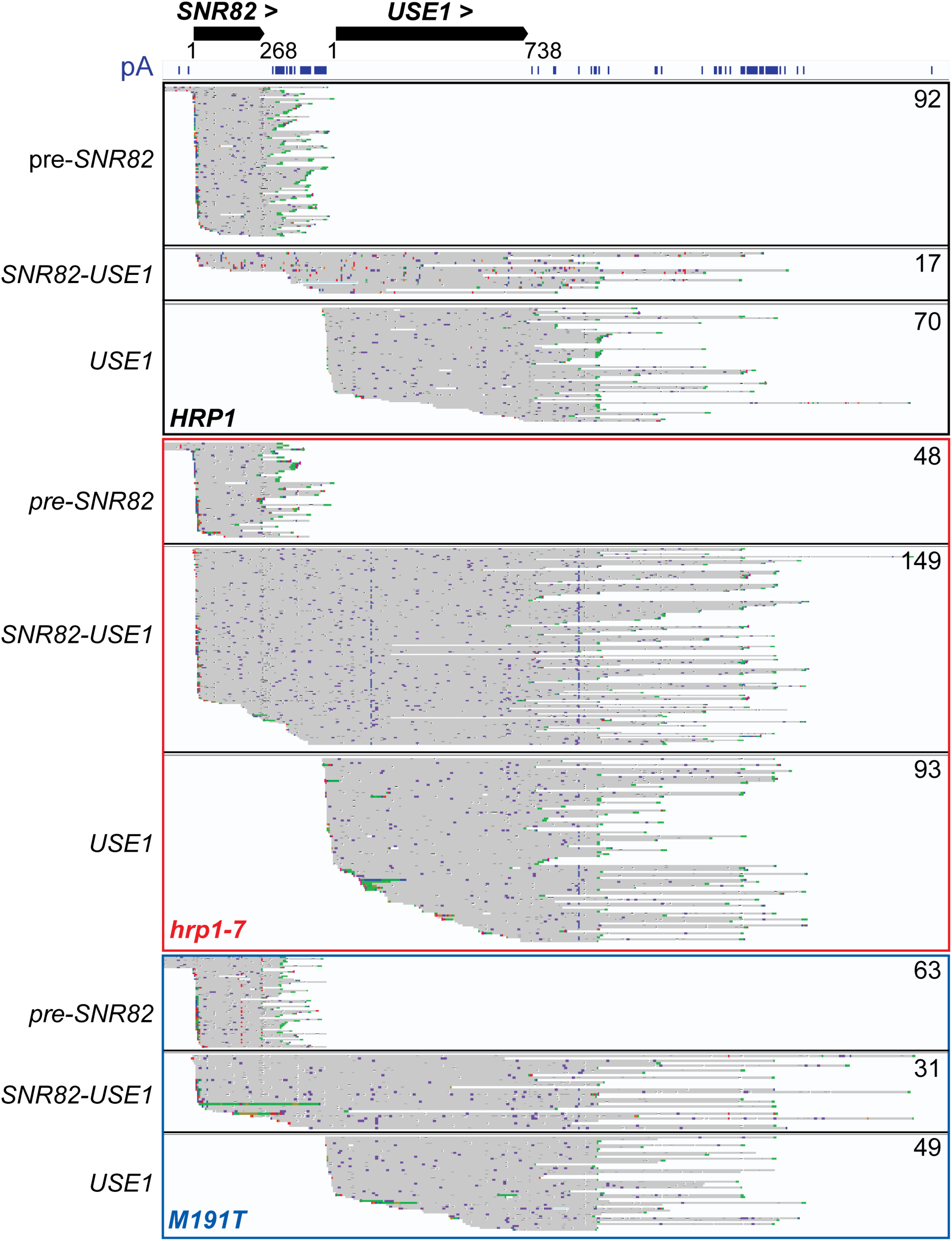
*hrp1-7* causes stronger readthrough of the *SNR82* terminator than *hrp1-M191T*. IGV browser view showing all dRNA-seq reads that intersect the *SNR82* and/or *USE1* loci, sorted by the regions spanned using NanoBlot (DeMario et al. 2023). The number of reads for each transcript class is shown in the upper right corner. Published polyA-seq data (pA) (Ozsolak et al. 2010) is shown below the ORF or mature snoRNA annotation. Reads from the *HRP1* strain are outlined in black, *hrp1-7* in red, and *hrp1-M191T* in blue. PolyA tails are marked in green and don’t represent the true length.

To examine the effects of *hrp1-7* on all sn/snoRNA terminators, we calculated the distance between the 3’ end of each read containing a sn/snoRNA and the annotated 3’ end of the mature snoRNA. All reads that end more than 500 nt downstream of a sn/snoRNA gene were classified as readthrough RNAs, which excluded most or all “properly” terminated pre-sn/snoRNAs. We then calculated the difference between the ratios of readthrough RNAs to total reads for each sn/snoRNA in *hrp1-7* and -M191T compared to *HRP1.* Eight out of 56 independently transcribed sn/snoRNAs have a greater than 10% average increase in readthrough in *hrp1-7* and 15 with greater than 5% average increase (Supplemental Table S3; Figure 3). Only *SNR82* and *SNR69* have a greater than 10% average increase in M191T compared to *HRP1* and six snoRNAs had at least 5% average increase. *SNR82* had the largest change with 7% readthrough in *HRP1*, 71% in *hrp1-7*, and 25% in M191T. Six of the 15 genes, *SNR33*, *SNR42*, *SNR69, SNR71, SNR80, and SNR82* were previously identified as being readthrough in the *hrp1-7* strain by microarray transcriptome analysis (Chen et al. 2017). Thus, only a small fraction of snoRNA terminators are sensitive to the mutations in *HRP1*. Perhaps the high transcription levels of snoRNA genes have necessitated the evolution of strong terminators with redundant elements.

**Figure 3.**
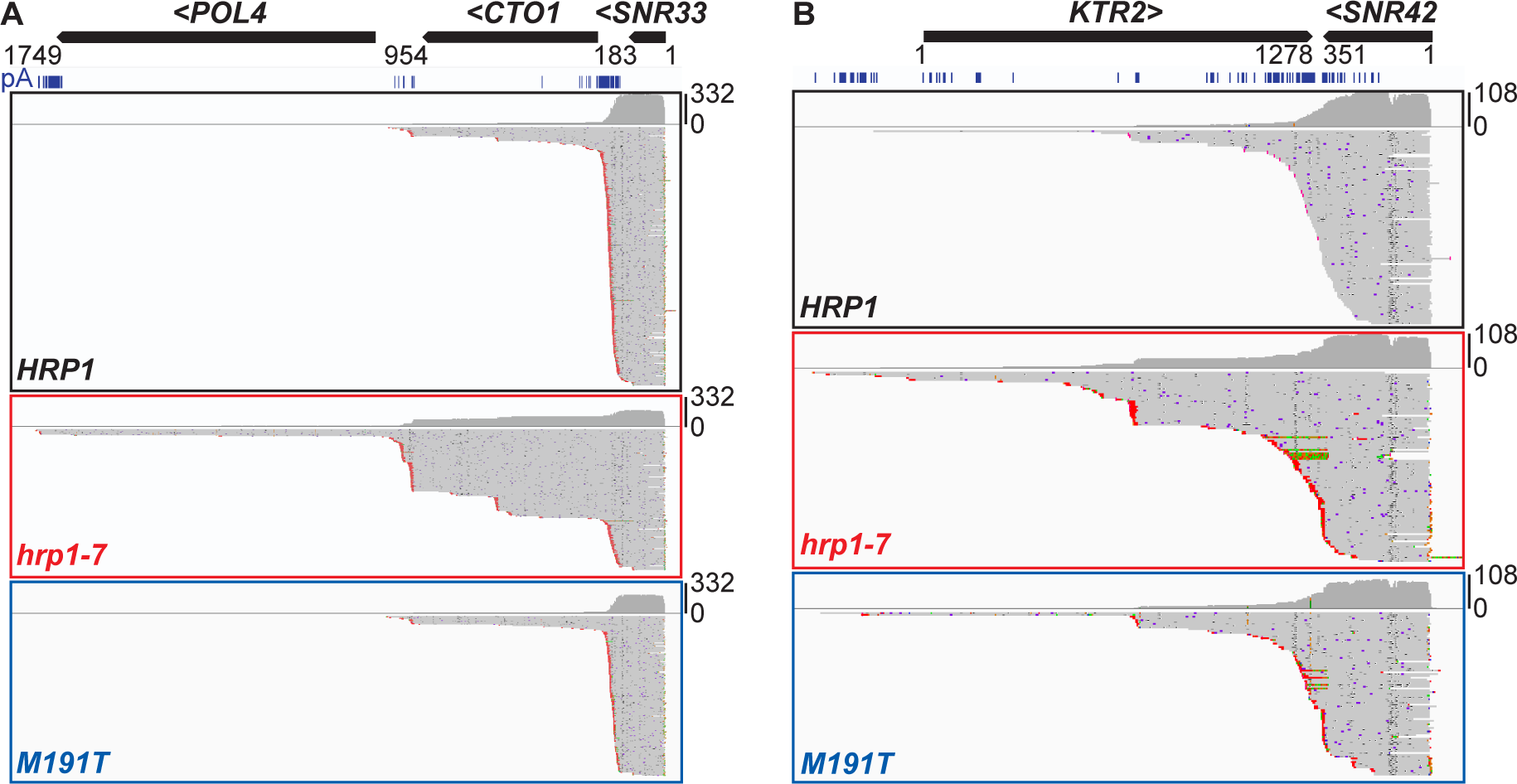
*hrp1-7* causes readthrough of *SNR33* and *SNR42* snoRNA terminators. IGV browser view showing all dRNA-seq reads that intersect snoRNA genes **A)** *SNR33* and **B)** *SNR42*. The data are displayed as in Figure 2, except the scale of the histogram is shown at far right. PolyA tails are marked in red for genes transcribed leftwards and don’t represent the true tail length.

A fraction of polyadenylated sn/snoRNA transcripts are likely degradation intermediates of mature sn/snoRNAs (Wyers et al. 2005). This is almost certainly true for U6 snRNA, encoded by the *SNR6* gene, which is made by RNAP III and not polyadenylated during synthesis. The polyadenylated U6 appears to have at least five nucleotides missing from the 5’ end, consistent with partial degradation (Supplemental Figure S2). A truncated form of the 1175 nucleotide U2 snRNA from the *LSR1* gene, which lacks the 5’ 12 nucleotides and ends 5-10 nucleotides downstream of the Sm-binding site for a total length of 110-120 nucleotides, is also likely a degradation intermediate. A similar sized 5’ U2 fragment was detected by Northern blot in several budding yeast species that have elongated U2 snRNAs (Roiha et al. 1989).

### Attenuated transcripts are readily detected from wild-type *HRP1*, *hrp1-7*, and *hrp1-M191T*

We next inspected protein-coding genes that contain known NNS attenuators, starting with *HRP1*. In the strains analyzed here, the genomic *HRP1* gene has its ORF replaced with the KanMX4 gene but the flanking sequences remain present. The intact *HRP1* allele encoding Hrp1 protein is carried on a centromere (low copy) plasmid. Since Hrp1 is essential, at least one copy of the plasmid must be present. Thus, the attenuated transcripts come from both the genome and the plasmid, while Hrp1 mRNA comes only from the plasmid and a Hrp1/KanMX fusion mRNA comes from the genome (Supplemental Figure S3A). With wild-type *HRP1*, there is a strong accumulation of attenuated reads whose 3’ ends cluster mainly upstream of or near the start codon of the *HRP1* ORF, consistent with a published polyA-seq dataset (Figure 4). The 3’ ends of these reads indicate that the *HRP1* attenuator contains two main regions that elicit termination, which we refer to as Term1 and Term2 (Supplemental Figure S3B). Strikingly, the total attenuated transcripts are about 6-fold greater than the full-length mRNA. From our earlier study we know that the *HRP1* attenuator is strong, since mRNA levels can increase more than 20-fold when productive translation of the mRNA is inhibited by mutation (Goguen and Brow 2023), but we did not expect that the attenuated transcripts would be stable enough to accumulate to such a high level. A potential caveat is that premature termination could be stronger on a plasmid-borne gene than on the chromosome. Also, mRNA containing the KanMX sequence may be less stable than intact Hrp1 mRNA, reducing the number of mRNA reads.

**Figure 4.**
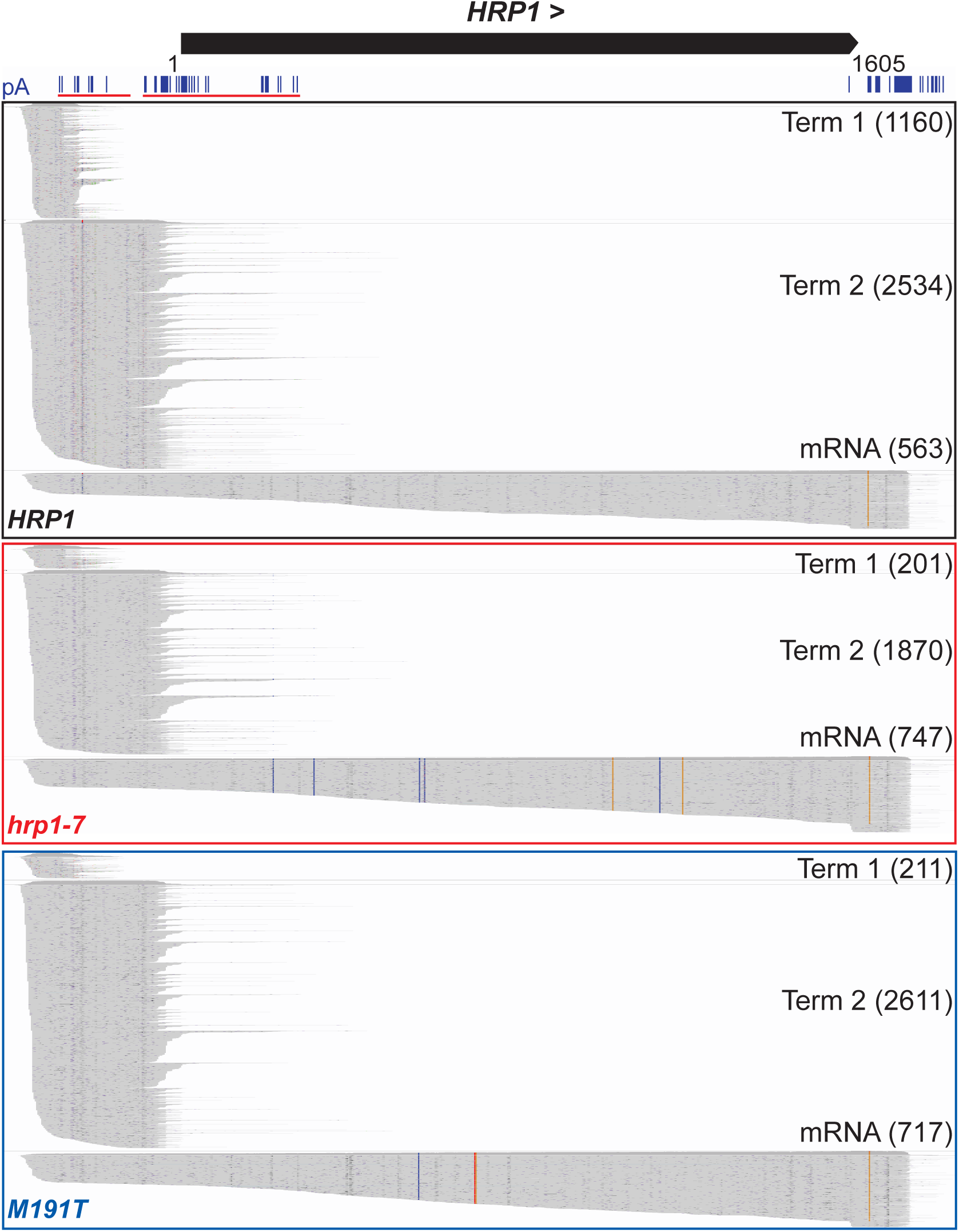
Attenuated *HRP1* RNAs accumulate in *HRP1*, *hrp1-7, and hrp1-M191T* strains. IGV browser view showing *HRP1* attenuated and full-length reads that have been sorted by the regions spanned using NanoBlot (DeMario et al. 2023). Published polyA-seq data (Ozsolak et al. 2010) is shown above (pA) with red underline marking Term1 and Term2 sites. Reads from the *HRP1* strain are outlined in black, *hrp1-7* in red, and M191T in blue with the number of reads for each transcript type shown in parentheses. The nucleotide substitutions in the *hrp1-7* and *-M191T* mRNA are confirmed by direct RNA-seq as indicated by the colored vertical lines, including a silent substitution in M191T to create a restriction enzyme site.

Finally, the cells were shifted to 37 °C for an hour before extracting the RNA to enhance the phenotype of *hrp1-7*. RNA from a strain that has only genomic *HRP1* and that was shifted to 18 °C for an hour before RNA extraction had fewer reads from attenuated transcripts than from Hrp1 mRNA (Wang et al. 2025).

A much smaller fraction of the attenuated transcripts end in Term1 in the *hrp1-7* and M191T strains than in the parental strain, consistent with increased readthrough of Term1 (Figure 4).

The total number of attenuated reads decreased 1.8-fold from wildtype to *hrp1-7*, consistent with readthrough of the dual attenuator, but the level of full-length mRNA did not increase correspondingly. If all the attenuated transcripts missing in the *hrp1-7* strain read through to the 3’-UTR, the mRNA should have increased 3.9-fold rather than the 1.3-fold increase observed, although half of this mRNA could contain KanMX4 sequence and been rapidly degraded, misaligned, or filtered out. Our previous results by Northern blot and RT-qPCR with probes in the Hrp1 ORF showed a four-to six-fold increase in *HRP1* mRNA in the *hrp1-7* strain (Kuehner and Brow, 2008; Goguen and Brow, 2023). The reason for this discrepancy is not clear, although it could be explained if most of the *hrp1-7* mRNA is not polyadenylated, as the prior analyses were done with unselected total RNA rather than polyA-selected RNA. In the *hrp1-M191T* strain, Term1 experienced similar readthrough as in the presence of Hrp1-7, but Term2 was less affected, resulting in only a 1.3-fold decrease in total attenuated transcripts.

Two other known attenuated genes, *NRD1* and *IMD2*, did not accumulate abundant truncated RNAs (Supplemental Figure S4), perhaps because the attenuated transcripts are less stable and/or these genes have weaker attenuators than *HRP1*. The ∼380-nucleotide *HRP1* 5’-UTR is highly conserved among the *Saccharomyces* genus so the attenuated transcripts are potentially functional. In contrast, the ∼290-nucleotide *NRD1* 5’-UTR is less well conserved.

### Identification of attenuators at genes that accumulate truncated RNAs

When we visually searched the transcriptome for other loci with high levels of 3’-truncated transcripts, the *PAP2/TRF4* gene was a striking example (Figure 5A). The truncated transcripts are approximately 330 bp long and end within the *PAP2/TRF4* ORF at codon 96 (Supplemental Figure S5). This 3’ end was also detected in wild-type cells previously (Ozsolak et al. 2010) indicating that these transcripts are present in other yeast strains. *hrp1-7* increases the accumulation of *PAP2/TRF4* truncated RNAs compared to the full-length roughly two-fold and *hrp1-M191T* has nearly as large an effect, indicating that formation of this 3’-end is influenced by Hrp1, directly or indirectly. Trf4 is a polyA polymerase that polyadenylates NNS-terminated transcripts (LaCava et al. 2005; Wyers et al. 2005; Ececioglu et al. 2006). A selection for increased readthrough of the *IMD2* attenuator yielded frame shift and nonsense mutations in *PAP2/TRF4* (Loya et al. 2012), as we have obtained in *NRD1* and *HRP1* in readthrough selections using other NNS terminators (Steinmetz and Brow 1996, Goguen and Brow 2023, Wang et al. 2025), suggesting that Trf4 may function in NNS termination as well as transcript processing and turnover. Accordingly, Trf4 could potentially autoregulate attenuation of its mRNA.

**Figure 5.**
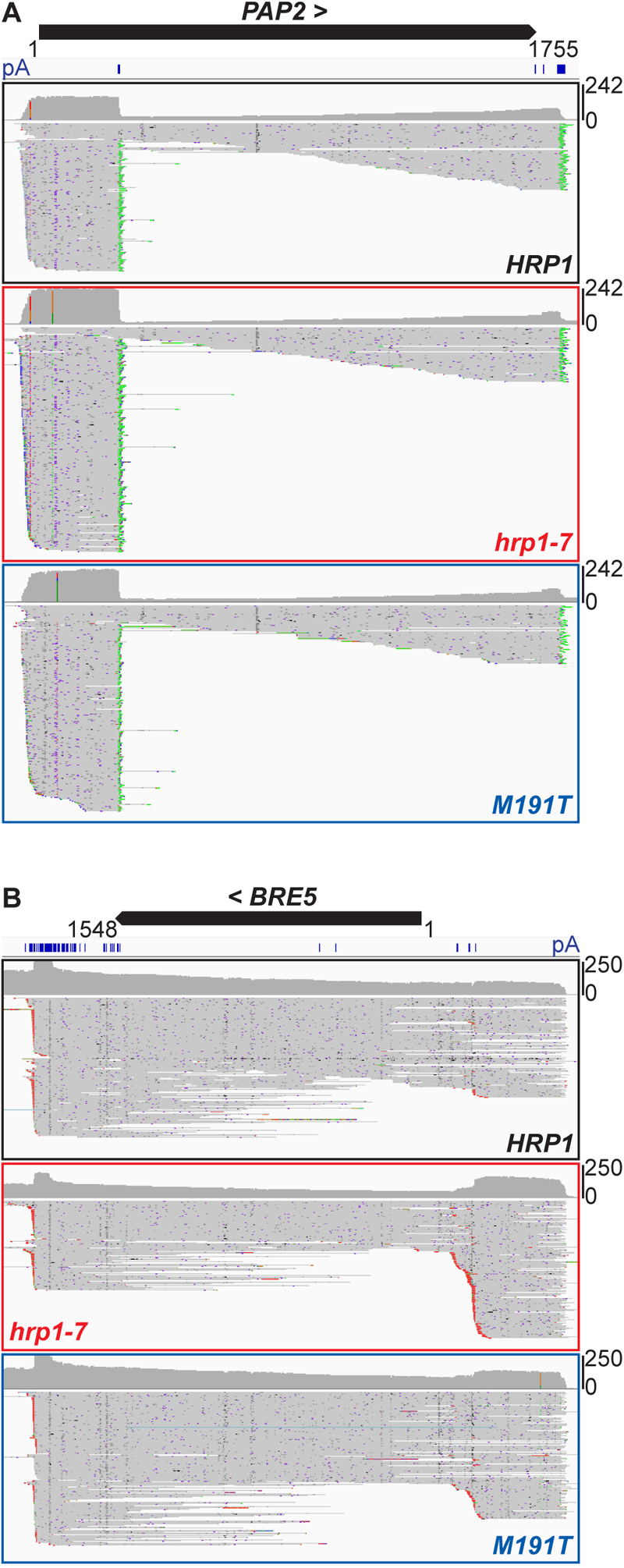
*HRP1* mutants increase 3’-end formation at potential novel attenuators. **A)** Truncated reads accumulate at the 5’ end of *PAP2/TRF4*. IGV browser view showing all reads that intersect with the *PAP2* locus. Published polyA-seq data (pA) (Ozsolak et al. 2010) is shown below the ORF. Reads from the *HRP1* strain are outlined in black, *hrp1-7* in red, and M191T in blue. **B)** IGV browser view showing all reads that intersect with the *BRE5* locus. *BRE5* is transcribed right to left.

We observed several other genes that accumulated truncated transcripts that end either before or within the 5’ end of the ORF. The levels of some of these potential attenuators were affected by *hrp1-7*, while others were not. *hrp1-7* causes a 3.5-fold accumulation of truncated RNAs in the 5’ UTR of the *BRE5* gene, indicating that it either increases recognition of this attenuator or increases the stability of these transcripts (Figure 5B). M191T shows a lesser accumulation of attenuated reads, indicating that the effect of *hrp1-7* requires more than one of the four substitutions.

### *hrp1-M191T* primarily induces readthrough of polyA sites, while *hrp1-7* additionally increases recognition of some ORF-internal sites

Short transcripts that terminate in the 5’-UTR of protein-coding genes, such as in *HRP1*, are classified as attenuated transcripts. When truncated transcripts include a start codon, as in the case of *PAP2/TRF4*, they could potentially be considered mRNA isoforms. When such transcripts exceed 500 nucleotides in length, they are more likely generated by CPA-dependent termination than NNS-dependent termination or may have “hybrid” terminators (Whalen et al. 2018; Amodeo et al. 2022). Previous studies mapping the 3’ ends of mRNAs have observed widespread ORF-internal polyadenylation (Ozsolak et al. 2010, Yoon and Brem 2010; Johnson et al. 2011, Pelechano et al. 2014). To better understand how these ORF-internal polyA sites are recognized we examined examples in *HRP1*, *hrp1-7*, and *hrp1-M191T* strains. We noted several genes in which use of ORF-internal polyA sites is increased relative to wild-type in the *hrp1-7* strain but not the *hrp1-M191T* strain (Figure 6).

**Figure 6.**
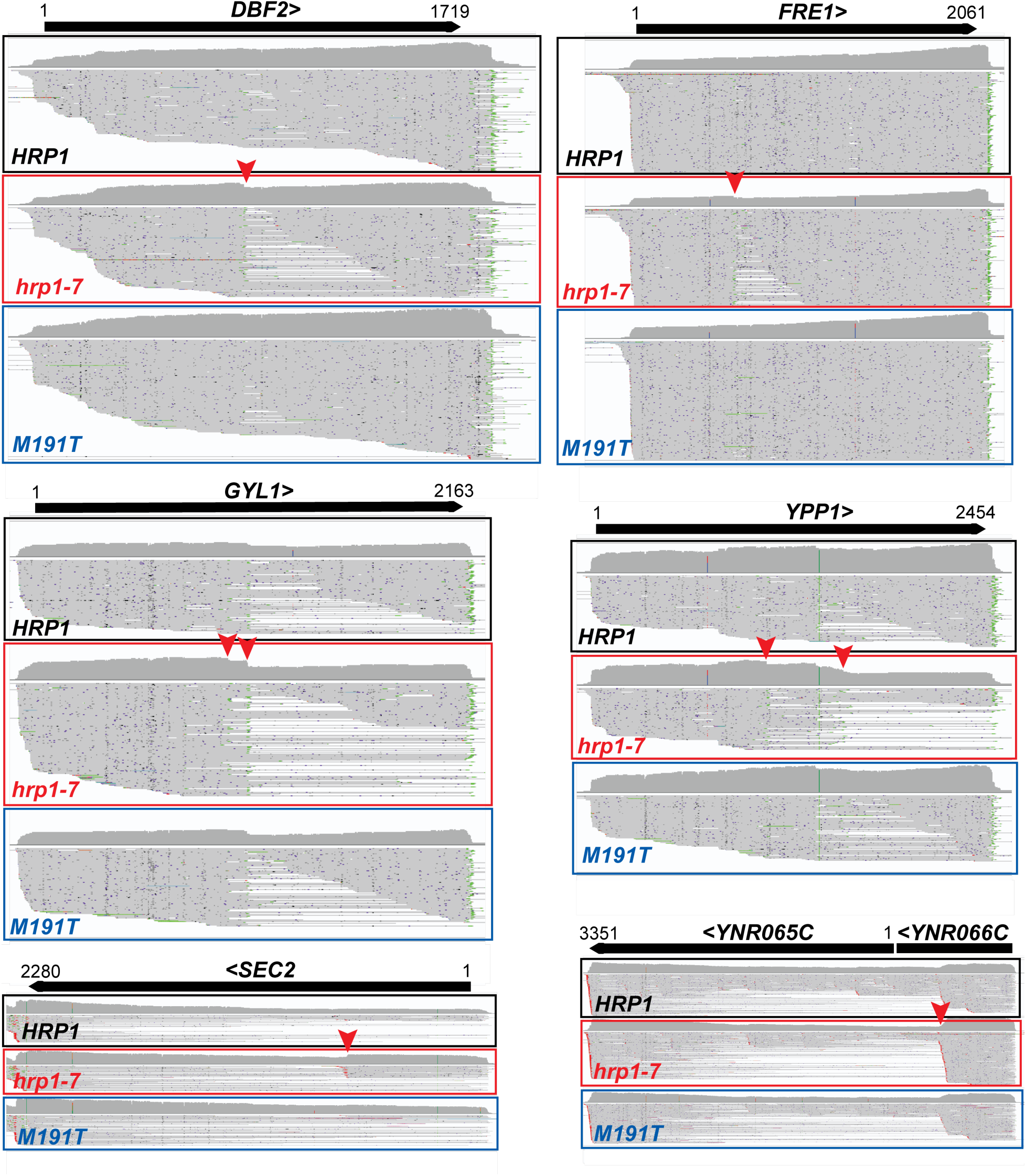
Examples of ORF-internal 3’-end formation increased by *hrp1-7* but not by *hrp1-M191T*. IGV browser view of dRNA-seq reads from the indicated genes in strains with the indicated *HRP1* genotype. Red arrowheads indicate 3’ ends that are more abundant in the *hrp1-7* strain than in the wild-type or *hrp1-M191T* strain. Not all reads are shown for *FRE1*. Genes in the bottom two panels are transcribed right to left. YNR066C and YNR065C appear to be contained in a single transcript.

To determine the net effect of each genotype on polyA site selection, we determined the distance of the polyA site in each read upstream (negative) or downstream (positive) of the stop codon for all protein-coding genes in the *HRP1, hrp1-7,* and *hrp1-M191T* strains, calculated the average polyA site position for each gene, and calculated the shift of the average position in each mutant strain (Figure 7A). We found that, globally, polyA sites are shifted downstream for both *hrp1-*7 and *hrp1-M191T* strains relative to the wild-type *HRP1* strain but the downstream shift is stronger for *hrp1-M191T* than *hrp1-7*. However, when looking at examples of mRNA 3’-UTRs, the downstream shifts caused by *hrp1-*M191T appear equivalent to or less than those caused by *hrp1-*7 (Figure 7B-D). Thus, the weaker global effect of *hrp1-*7 on polyA site readthrough is likely due to a propensity to cause premature termination in the ORF, which suggests the possibility that one or more of the other three substitutions in *hrp1-7* may promote ORF-internal polyA site recognition. This behavior of *hrp1-7* is potentially due to a reduced RNAP II elongation rate, which has been shown to cause an upstream shift in polyA site profiles (Geisberg et al. 2020) (see Discussion).

**Figure 7.**
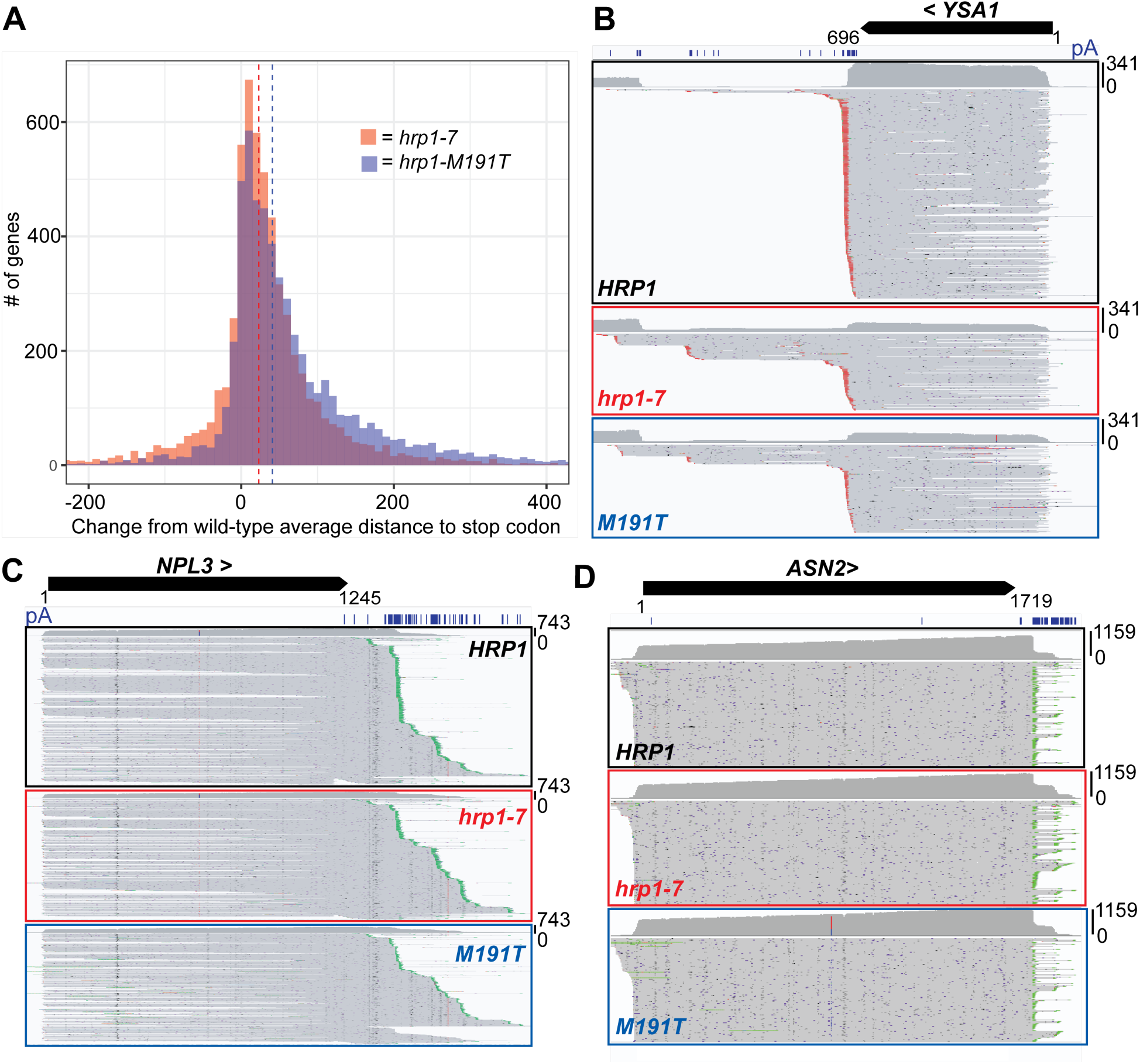
The stronger average downstream shift of polyA sites in *hrp1-*M191T than *hrp1-7* is not due to 3’-UTR polyA sites. **A)** Histogram (bin width = 10) of the shift in average polyA site position relative to the stop codon in *hrp1* mutant strains relative to wild-type. Dotted lines show the median values for *hrp1-7* (red) and *hrp1-M191T* (blue). **B-D)** IGV browser view showing reads that intersect with the indicated genes. *hrp1-M191T* exhibits a weaker or similar downstream shift of polyA sites compared to *hrp1-7*. Not all individual reads are shown for *ASN2*.

### *YRA1* and *DBP2* mRNA are decreased in *hrp1-7* but not *hrp1-M191T* strains

Yra1 is an mRNA-binding protein that has been implicated in polyA site selection (Johnson et al. 2011) and is negatively autoregulated by the presence of an unusually large intron (Rodríguez-Navarro et al. 2002; Preker et al. 2002). Its mRNA level is noticeably decreased in the *hrp1-7* strain, but not the *hrp1-M191T* strain, which may have a small increase in unspliced Yra1 mRNA (Figure 8A). This result indicates the other *hrp1-7* substitutions contribute to the change in steady state level of Yra1 mRNA. Dbp2 is a DEAD-box helicase that physically and functionally interacts with Yra1 (Ma et al. 2013) and its level is also negatively autoregulated by a large intron (Barta and Iggo 1995). Dbp2 mRNA levels also decrease in the *hrp1-7* strain and, conversely, increase in the *hrp1-M191T* strain, without any noticeable change in splicing efficiency (Figure 8B).

**Figure 8.**
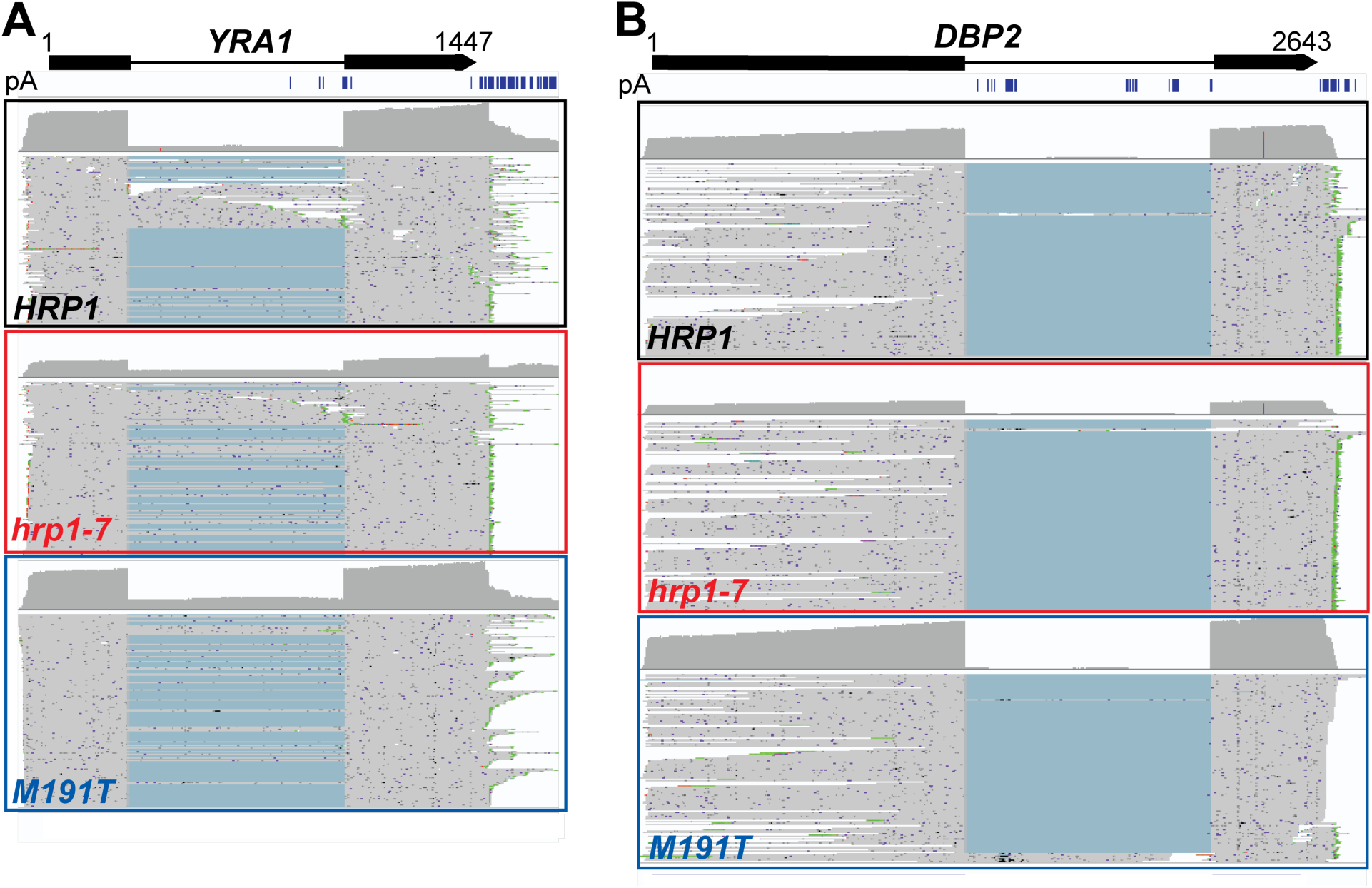
Decreased Yra1 and Dbp2 mRNA in *hrp1-7*. IGV browser view of dRNA-seq reads from the indicated genes in strains with the indicated *HRP1* genotype. Summed reads are shown in dark gray above each read stack. Within each panel, all summed reads are to the same scale: **A)** *YRA1* = 0 to 684 reads, **B)** *DBP2* = 0 to 1386 reads. Blue lines connect spliced exons. Not all individual reads are shown. Published polyA-seq data (pA) (Ozsolak et al. 2010) is shown below the ORF and intron.

The mechanism of intron-dependent autoregulation is not fully understood for either *YRA1* or *DBP2* (Preker and Guthrie 2006; Dong et al. 2007). The presence of polyA sites in both introns and our observation that Hrp1-7 results in lower mRNA levels from both genes (Figure 8) suggests that mRNA downregulation is due to intronic CPA and rapid degradation of the 3’-truncated pre-mRNA. The increased utilization of ORF-internal polyA sites seen in the presence of Hrp1-7 (Figure 6) could also occur in these two introns. This mechanism can explain the increase in *DBP2* mRNA in the presence of Hrp1-M191T, which may cause readthrough of the intronic polyA sites. It also can explain the position and length dependence of the introns for autoregulation, as the polyA sites must be far enough downstream of the transcription start site to be efficiently recognized by the CPA factors. This model implies that Yra1 and Dbp2 promote recognition of the intronic polyA sites to downregulate their respective mRNAs.

## DISCUSSION

### Nanopore direct RNA-seq efficiently detects altered 3’-end formation

Both NNS- and CPA-termination are coupled to polyadenylation of the nascent transcript, by the TRAMP and CPF complexes, respectively. Thus, both can be visualized by dRNA-seq using an oligo(dT) splint to selectively target ligation of the sequencing adapter to these transcripts.

No oligo(dT)-affinity purification or rRNA depletion is necessary and one microgram of yeast total cellular RNA resulted in an average transcriptome read depth of at least 175 in 24 hours (Supplemental Table S1). Reads from mature non-coding RNAs are greatly reduced as they are not polyadenylated except during degradation. Longer reads are 3’-biased since the 3’-end of each RNA is inserted into the sequencing pore first, but for most genes enough full-length reads are obtained to accurately map 5’-ends.

The large number of attenuated transcripts we observed for the *HRP1* gene is consistent with the large dynamic range of Hrp1 mRNA expression we previously found in the presence of mutations. For example, substitution of the AUG start codon to ACG or GUG results in a 25-to 40-fold increase in mRNA as measured by qRT-PCR (Goguen and Brow 2023). The wild-type attenuated transcripts were not detected in our prior microarray transcriptome analysis (Chen et al. 2017), perhaps in part due to missing (non-unique) probes and smoothing of the results.

Conversely, the ∼5-fold increase in Hrp1 mRNA in the *hrp1-7* strain after temperature shift measured by Northern blot (Kuehner and Brow 2008) was not seen in the dRNA-seq data, which detected only a 1.2-fold increase. More experience with dRNA-seq will be required to fully understand its strengths and weaknesses.

The accumulation of transcripts terminating upstream of the start codon of protein-coding genes identified several new attenuated genes, most notably *PAP2/TRF4*, but also *ACC1, BRE5, GCN4, MAL31, PHO80, SEN1, SFP1, TPO4,* and *YOL036W*. This method of detecting attenuation requires that the attenuated transcripts accumulate at steady state, due to sufficient synthesis and stability. Therefore, some attenuated genes may be missed by this method. The accumulation of 3’-ends within protein-coding regions may be cases of NNS-dependent attenuation, internal cleavage and polyadenylation, or some hybrid of the two pathways. There is no way to discriminate between these possibilities from our data. However, for many examples the 3’-end is more than 800 nucleotides from the cap site, including *BOP2, CHS6*, *DBF2, DIA2, DSS1, GYL1, MIT1, PEX27, PRP8,* and *RNA14*. These 3’-ends are outside the range where NNS termination is expected to occur, typically 500 nucleotides from the cap site or less. We noted one case of an intronic polyA site, in intron 2 of the *EFM5* gene. Use of this site was increased in *hrp1-7* but not *hrp1-M191T* and resulted in a deletion of exon 3 from some mRNAs from this gene.

### Evidence that Hrp1 is an anti-termination factor

The cleavage and polyadenylation efficiency element, (UA)_3_, binds Hrp1 with high affinity and has long been recognized as important for mRNA 3’-end formation in yeast cells (Guo and Sherman 1996, Kessler et al. 1997, Chen and Hyman 1998, Minvielle-Sebastia et al. 1998).

Recently, a selection for sequences in a random 50-mer oligonucleotide that increased expression of a yeast reporter mRNA when present in the 3’-UTR yielded (UA)_3_ as the top hexamer (Savinov et al. 2021). But the mechanism by which Hrp1 binding enhances mRNA synthesis, stability, and/or protein expression is not known. Our finding that Hrp1 promotes NNS termination of some genes suggests that it does not act solely by interaction with CPA factors. Our recently identified genetic interaction between Hrp1 and the Rpb3 subunit of RNA polymerase (Wang et al. 2025) supports a model whereby Hrp1 promotes termination of RNA synthesis in both the CPA and NNS pathways by slowing RNAP II elongation upon binding to (UA)_3_ in the nascent transcript (Goguen and Brow 2023).

In this study, our finding that the *hrp1-7* mutant results in either upstream or downstream shifts in polyA sites to roughly similar extents (Figure 7A) can be explained by the four substitutions altering two different activities of Hrp1: sequence-specific binding to RNA and interaction with the RNAP II elongation complex (EC). Substitutions that weaken RNA-binding are expected to result in a downstream shift in polyA sites and readthrough of Hrp1-dependent NNS terminators, while substitutions that weaken binding to and/or allosteric activation of the RNAP II EC are expected to result in an upstream shift in polyA sites and reduced readthrough of Hrp1-dependent NNS terminators. This interpretation is in keeping with the “kinetic coupling” model of 3’-end formation, which states that the probability of recognition of a terminator increases with a decreased rate of RNAP II elongation (see Kuldell and Kaplan 2025 for a recent discussion).

Our finding that the Hrp1-M191T substitution decreases the affinity for (UA)_4_ in vitro and results primarily in readthrough of NNS terminators and CPA sites supports the hypothesis that Hrp1 has RNAP II antitermination activity that is diminished upon specific binding to the nascent transcript. However, snoRNA terminator readthrough induced by Hrp1-M191T was generally weaker than that conferred by Hrp1-7, suggesting that one or more of the other three substitutions contribute to, rather than antagonize readthrough. These substitutions are: i) in RRM1 near M191T (A195P), in an Asn-and Gly-rich stretch between RRM2 and the Met/Gln-rich low complexity domain (N345D), and within the Met/Gln LCD (Y383H). Hrp1-N345D alone induced 1.8-fold readthrough of the *HRP1* attenuator and may synergize with M191T (3-fold readthrough) to produce the 8-fold readthrough effect of *hrp1-7* on its attenuator (Goguen and Brow 2023). Neither Hrp1-A195P nor -Y383H had any effect on *HRP1* attenuator readthrough, and thus are candidates for eliciting early termination, for example, by weakening association with the RNAP II elongation complex (EC). Hrp1-A195P reproducibly decreased expression of an *HRP1-CUP1* reporter as judged by increased copper sensitivity, potentially due to increased premature termination (Goguen and Brow 2023). Y383H and three other substitutions in the Met/Gln-rich LCD partially suppress the cold-sensitivity of the Rpb3-K9E substitution in RNAP II, potentially by weakening binding to the RNAP II EC (Wang et al. 2025). Together, these two substitutions could be responsible for increased use of ORF-internal polyA sites in some genes in the presence of Hrp1-7. However, we cannot rule out that the increased recognition of some ORF-internal polyA sites by Hrp1-7 is due to a change in specificity of RNA-binding.

Our dRNA-seq data does not clearly reproduce the global decline in RNA abundance along the length of protein coding genes that we observed in our microarray transcriptome data of total RNA from an *hrp1-7* strain, which first suggested a function of Hrp1 in antitermination (Chen et al. 2017). This discrepancy could, at least in part, be due to the failure of some reads to reach the 5’ ends of mRNAs, which would tend to mask a decrease in transcript abundance along genes. It could also be explained if the 3’-truncated transcripts are not efficiently polyadenylated and thus are not detected by our dRNA-seq protocol, although it is not obvious how this might occur. In any case, as discussed above, several other lines of evidence point to a function of Hrp1 in antitermination.

### Possible pleiotropic effects of substitutions in Hrp1

Because Hrp1 functions in both co- and posttranscriptional processes and its binding sites remain in the processed mRNA, substitutions in Hrp1 could have effects on mRNA biosynthesis, transport, translation, and turnover. Since dRNA-seq detects steady-state RNAs, both synthesis and degradation (including deadenylation) can contribute to changes in RNA levels and length. In addition to the possibility of indirect effects on target genes, for example, changes in levels of relevant transcription activators, numerous downstream processes could be altered. A significant challenge in understanding the effects of sequence variants on the function of nuclear RNA-binding proteins will be dissecting out the influence of the variants on different steps of the RNA’s lifecycle.

## MATERIALS AND METHODS

### Plasmids

pET41a-HRP1 is derived from pDL469, containing a C-terminal TEV protease cleavage peptide (ENLYFQG) followed by a 6x His-tag (Tudek et al. 2014). The coding region of Nab3 in pDL469 was replaced with the coding region of Hrp1 from pRS313-HRP1 (Goguen and Brow 2023) using NEB Builder DNA HiFi Assembly. To construct pET41a-hrp1-7 and pET41a-hrp1-M191T, pRS313-hrp1-7 and pRS313-hrp1-M191T (Goguen and Brow, 2023) were digested with SacI and AatII and gel-purified using the GeneJET Gel Extraction Kit (Thermo Scientific). The fragments were then ligated with the SacI/AatII digested and gel-purified backbone of pET41a-HRP1.

### Recombinant protein expression and purification

pET41a plasmids were transformed into Rosetta2 DE3 pLysS cells (Novagen). Cells were grown to OD_600_ = 0.6-0.8 in 1 L of Terrific Broth media (Fisher) at 37°C. Protein expression was induced by adding 0.1 mM isopropyl-D-1-thiogalactopyranoside. The induced culture was incubated at 16°C for 16-20 hours, then harvested by centrifugation. Cell pellets were resuspended in 50 mL of IMAC buffer (50 mM sodium phosphate, 500 mM NaCl, 5% glycerol, 1 mM DTT, pH 7.5) + 20 mM imidazole and 1x EDTA-free protease inhibitor cocktail (Calbiochem). Cells were lysed by sonication (D100, Fisher Scientific) at 50% power for 20 cycles of 10 seconds of sonication with 30 second rests on ice, followed by centrifugation at 20,000 x *g* for 15 minutes and 0.45 um filtration of the supernatant. Five mL Ni-NTA resin (Qiagen) pre-equilibrated with IMAC buffer + 20 mM imidazole was added directly to filtered supernatant and incubated with rocking for 1 hour at 4°C. Lysate and resin were added to a glass column, washed with 5 column volumes (CV) of IMAC buffer + 20 mM imidazole, washed with 5 CV of IMAC buffer + 50 mM imidazole, then eluted in 3 CV of IMAC buffer + 300 mM imidazole. Protein-containing fractions based on OD_280_ absorbances and SDS-PAGE analysis were pooled, 2 mg of His6-tagged TEV protease was added, and the mixture dialyzed into buffer containing 50 mM Tris, 250 mM NaCl, 20 mM imidazole, 1 mM DTT, 5% glycerol using Thermo Scientific Slide-A-Lyzer Dialysis Cassette (10 kD MWCO, 3-12 mL capacity). Cleavage of the 6x His tag from the Hrp1 constructs proceeded overnight at 4°C. The volume of the sample was brought to 5 mL by concentration using Amicon centrifugal filter unit 10 kDa MWCO (Millipore) and run through a 1 mL IMAC HiTrap HP column (Cytiva) on the AKTA pure FPLC system. The flowthrough of the column was collected, concentrated to 0.5 mL as before, and further purified on a Superdex 200 Increase 10/300 GL column (Cytiva) in buffer containing 50 mM Tris pH 7.5, 500 mM NaCl, 10% glycerol, and 1 mM DTT. Peak fractions at OD_280_ were run on a 4-15% Mini-PROTEAN TGX Pre-cast Gel (Bio-Rad) and stained using Thermo Scientific Silver Stain kit to assess purity. Pooled fractions were buffer exchanged into storage buffer (20 mM Tris-HCl pH 8, 150 mM NaCl, 1 mM DTT, 20% glycerol) using Amicon spin filters and stored in aliquots at -80°C.

### RNA-binding assay

Fluorescence anisotropy experiments were conducted in a Tecan Spark microplate reader. Samples were excited vertically at 490 nm and horizontal and vertical emission at 520 were recorded. The anisotropy values were automatically measured by the Fluorescence Standard Polarization function of SparkControl software. All measurements were conducted at 25°C in buffer containing 20 mM Tris pH 8, 150 mM NaCl, 5% glycerol, and 1 mM DTT. 20 nM of FAM-labeled (UA)_4_ RNA (Sigma) was added to increasing concentrations of full-length Hrp1 or mutant protein in a black flat-bottom 96-well plate (Greiner) on ice. Anisotropy measurements were taken every 5 minutes after the plate reader reached ambient temperature for 30 minutes total. Binding curves were constructed using the anisotropy values from the 15-minute timepoint. The experimental binding isotherms were analyzed using one-site specific binding equation in GraphPad Prism. B_max_ for all three proteins was constrained to the value for the wild-type protein.

### Yeast strains

The yeast strains used in this study are derived from ECG004 (*MATalpha hrp1τ<::KanMX4 cup1τ<::ura3 ura3-52 his3-τ<200 trp1-τ<63 lys2-801 ade2-1 leu2-τ<1* [pRS316-HRP1]) (Goguen and Brow, 2023). Strains ECG012, ECG013, and ECG014 were made by replacing pRS316-HRP1 with pRS313-HRP1, -hrp1-7, or -hrp-M191T, respectively (Goguen and Brow, 2023).

### RNA purification

Yeast cells were grown in YEPD at 30°C to an OD_600_ of 0.6-1.0, quickly shifted to 37°C by adding an equal volume of 44°C YEPD, then incubated for 1 hr at 37°C, and pelleted by centrifugation. Total cellular RNA was prepared with the GeneJet RNA Purification Kit (Thermo Scientific) according to the manufacturer following cell lysis by vortexing with glass beads (200-325 um diameter, Sigma-Aldrich) in the kit Lysis Buffer supplemented with 40 mM DTT. Total RNA was treated with RNase-free DNase I (NEB) followed by purification with the GeneJET RNA Purification Kit using the RNA clean-up protocol. RNA concentrations were measured using a Qubit fluorometer and Qubit RNA Broad Range assay kit (Invitrogen). RNA integrity was determined using the Agilent RNA ScreenTape and TapeStation 4150 system. All samples used for library preparation had an RNA Integrity Number equivalent (RINe) score of 10.

### Nanopore library preparation and sequencing

Direct RNA-seq libraries were prepared with 1 μg of total cellular RNA using the Oxford Nanopore Direct RNA sequencing kit SQK-RNA004 (ONT, protocol version Sept. 20, 2023) following the manufacturer’s instructions except that SuperScript IV Reverse Transcriptase (Thermo Fisher Scientific) was used instead of SuperScript III Reverse Transcriptase. Sequencing was performed using FLO-MIN004RA flow cells on a MinION Mk1B device.

Raw voltage traces were collected as POD5 files using ONT MinKnow software (versions 5.8.7 and 6.2.6). POD5 files were base-called using Dorado (v0.7.2) high accuracy base-calling (hac) with --estimate-poly-a parameter. Base-called reads were aligned to the *Saccharomyces cerevisiae* genome (S288C, RefSeq R64, GCF_000146045.2) using the EPI2ME Alignment workflow (ONT, v1.2.0) with the minimap2 preset override: -ax splice -uf -k14 -G2k. Maximum intron size (-G) was set to 2 kb to prevent misalignment by the addition of large introns. See Supplemental Table S1 for basecalling and alignment summaries. Alignments were visualized with the Integrated Genome Viewer (IGV) (Robinson et al. 2011).

### Data analysis

All data analysis was conducted using RStudio (v4.3.3) and graphs were produced using ggplot2 (v3.5.0). Genomic coordinates for all annotated genes were obtained from the Saccharomyces Genome Database (SGD) All Annotated Sequence Features (Engel et al. 2025). Reads were assigned to genes using genomicRanges (v1.54.1) findOverlaps function with a required minimum overlap (minoverlap) of 50 bp to prevent misassignment of nearby genes on the same strand. Reads were filtered for a minimum map quality score of 20. Read counts per gene were normalized to overall read depth for each sample to produces RPM (Reads Per gene per Million) for correlation analysis. Visualization of reads intersecting individual genes were isolated using NanoBlot (DeMario et al. 2023) subsetNanoblot function. ORF and 3’ UTR polyA sites were classified based on their position relative to the annotated stop codon of protein coding genes (Engel et al. 2025).

## ACKNOWLEDGEMENTS

We thank Odil Porrua and Domenico Libri for plasmid pDL469, Yuichiro Nomura and Sam Butcher for the plasmid and protocol for expressing and purifying TEV protease, and Moyao Wang for discussions. This work was supported by NIH grant R35 GM118075 to DAB and funding from the Department of Biomolecular Chemistry for ECG. Fluorescence anisotropy data were acquired at the UW-Madison Biophysics Instrumentation Facility, which was established with support from UW-Madison and grants BIR-9512577 (NSF) and S10 RR13790 (NIH).

