## Supplemental tables and figures for "Direct RNA-seq provides evidence for an antiterminator function of the yeast Hrp1 protein on RNA polymerase II transcripts"

**Supplemental Table S1.** Nanopore sequencing and alignment.

| Strain: | Flow cell run time: | Dorado base-called reads: | Minimap2 aligned reads: | Total bases called and mapped: |
|---|---|---|---|---|
| HRP1 rep 1 | 23.0 hrs | 2.72 M | 2.56 M | 2.27 GB |
| hrp1-7 rep 1 | 23.0 hrs | 2.46 M | 2.40 M | 2.41 GB |
| HRP1 rep 3 | 24.0 hrs | 3.71 M | 3.28 M | 3.38 GB |
| hrp1-7 rep 3 | 25.5 hrs | 2.37 M | 1.93 M | 2.02 GB |
| hrp1-M191T | 27.0 hrs | 2.70 M | 2.61 M | 2.58 GB |

**Supplemental Table S2.** Pairwise Spearman correlation coefficients between *HRP1* and *hrp1-7* biological replicates.

| | HRP1 rep 1 | HRP1 rep 3 | hrp1-7 rep 1 | hrp1-7 rep 3 |
|---|---|---|---|---|
| HRP1 rep 1 | | | | |
| HRP1 rep 3 | 0.9870 | | | |
| hrp1-7 rep 1 | 0.9743 | 0.9760 | | |
| hrp1-7 rep 3 | 0.9686 | 0.9742 | 0.9849 | |

**Supplemental Table S3.** Difference in % readthrough of independently transcribed snoRNA genes in *hrp1-7* and *hrp1-M191T* strains relative to the parental *HRP1* strain. SnoRNAs with 10% or greater increase in readthrough are shown in red and 5% or greater shown in orange.

| Gene name | Avg hrp1-7 - HRP1 | Avg M191T - HRP1 |
|---|---|---|
| SNR82 | 0.64 | 0.18 |
| SNR33 | 0.49 | 0.00 |
| SNR64 | 0.35 | 0.05 |
| SNR161 | 0.24 | -0.01 |
| SNR69 | 0.21 | 0.11 |
| SNR68 | 0.19 | 0.03 |
| SNR42 | 0.15 | 0.05 |
| SNR5 | 0.11 | 0.01 |
| SNR58 | 0.09 | 0.00 |
| SNR80 | 0.09 | 0.05 |
| SNR71 | 0.06 | 0.03 |
| SNR9 | 0.06 | 0.05 |
| SNR46 | 0.06 | 0.01 |
| SNR79 | 0.05 | 0.09 |
| SNR8 | 0.05 | 0.02 |
| SNR50 | 0.03 | 0.00 |
| SNR60 | 0.03 | -0.02 |
| SNR85 | 0.02 | -0.01 |
| SNR39B | 0.02 | 0.00 |
| SNR52 | 0.02 | 0.02 |
| SNR13 | 0.01 | -0.03 |
| SNR49 | 0.01 | 0.00 |
| SNR30 | 0.01 | 0.01 |
| SNR10 | 0.01 | 0.00 |
| SNR34 | 0.01 | 0.04 |
| SNR11 | 0.00 | 0.03 |
| SNR6 | 0.00 | 0.01 |
| SNR63 | 0.00 | 0.00 |
| SNR86 | 0.00 | 0.00 |
| SNR14 | 0.00 | 0.00 |
| SNR17A | 0.00 | 0.00 |
| SNR17B | 0.00 | 0.00 |
| SNR19 | 0.00 | 0.00 |
| SNR32 | 0.00 | 0.00 |
| SNR36 | 0.00 | 0.00 |
| SNR40 | 0.00 | 0.00 |
| SNR56 | 0.00 | 0.00 |

| | | |
|---|---|---|
| SNR65 | 0.00 | 0.00 |
| SNR83 | 0.00 | 0.00 |
| SNR4 | 0.00 | 0.00 |
| SNR45 | 0.00 | -0.01 |
| SNR31 | 0.00 | 0.00 |
| SNR84 | 0.00 | 0.00 |
| SNR43 | -0.01 | -0.01 |
| SNR35 | -0.01 | -0.01 |
| SNR62 | -0.01 | 0.00 |
| SNR47 | -0.01 | -0.01 |
| SNR81 | -0.01 | 0.01 |
| SNR37 | -0.01 | -0.02 |
| SNR66 | -0.01 | -0.01 |
| SNR7-L | -0.02 | -0.02 |
| SNR7-S | -0.02 | -0.02 |
| SNR48 | -0.02 | 0.00 |
| SNR189 | -0.04 | -0.07 |
| SNR87 | -0.08 | 0.03 |
| SNR3 | -0.09 | -0.08 |

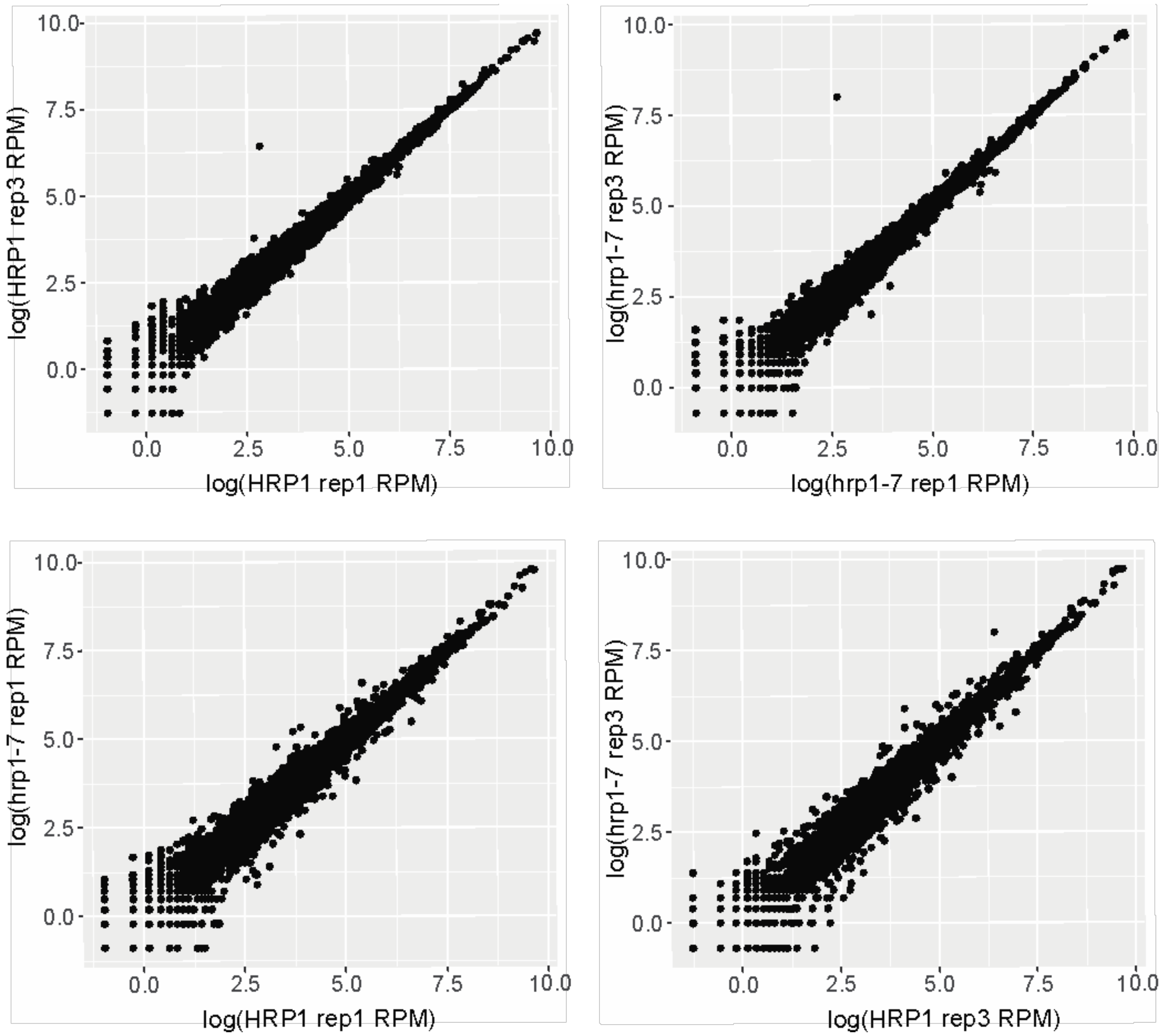


**Supplemental Figure S1.** Scatterplots showing the correlation between of normalized reads per million (RPM) per gene between biological replicates of *HRP1* and *hrp1-7* strains.

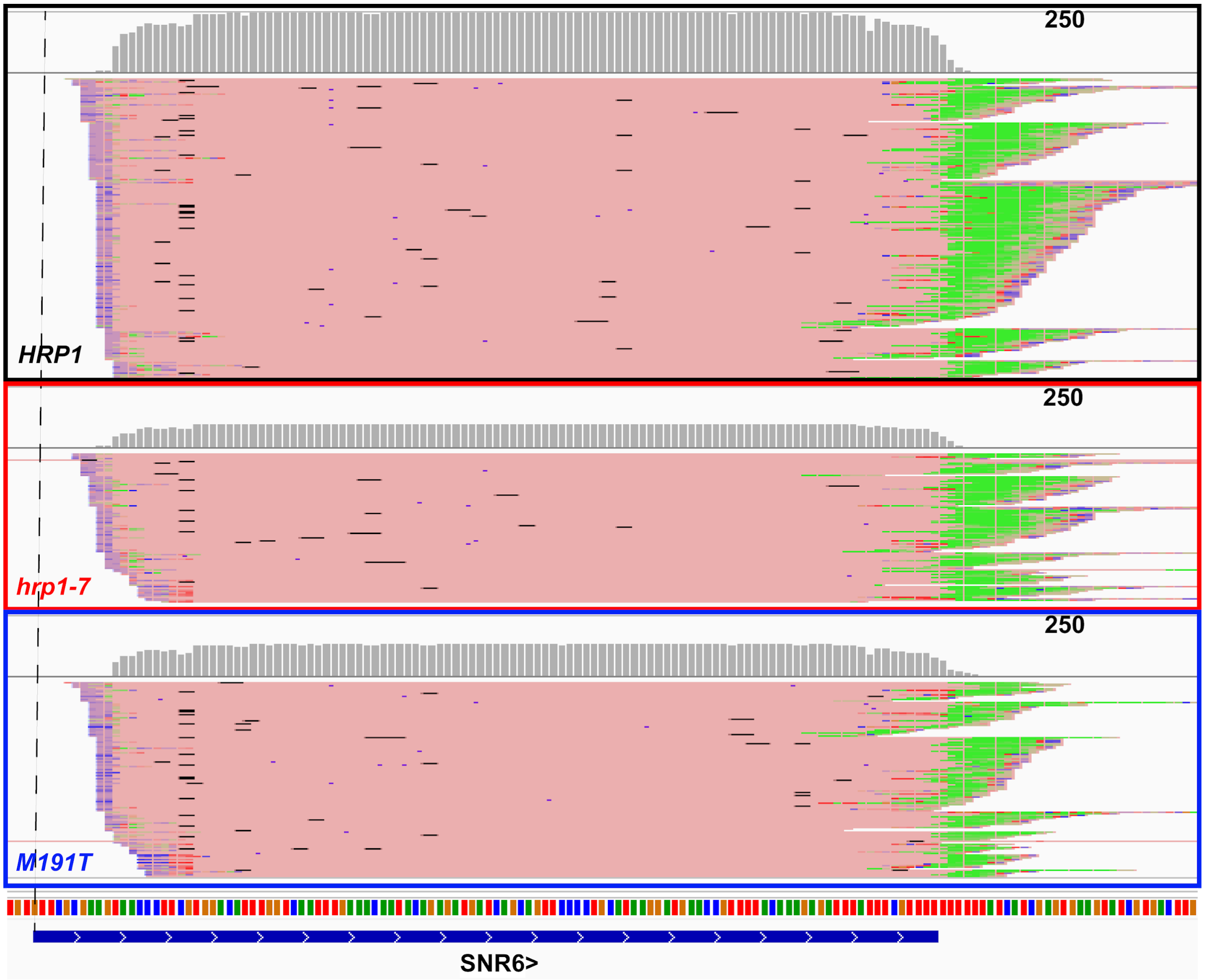


**Supplemental Figure S2.** IGV browser view showing dRNA-seq reads aligned to the U6 snRNA gene, *SNR6*, in isogenic strains with the indicated *HRP1* genotype. Not all indivdual reads are shown. The blue bar at the bottom shows the extent of mature U6 snRNA, the colored marks above it represent the nucleotide sequence, and the dotted line marks the first nucleotide of mature U6. The polyA tail is colored green. The gray histogram at the top of each panel shows the number of reads for each nucleotide. The scale is 0-250 reads for all three panels. The reason for the decreased abundance of the polyadenylated U6 snRNA in the mutant strains is not known.

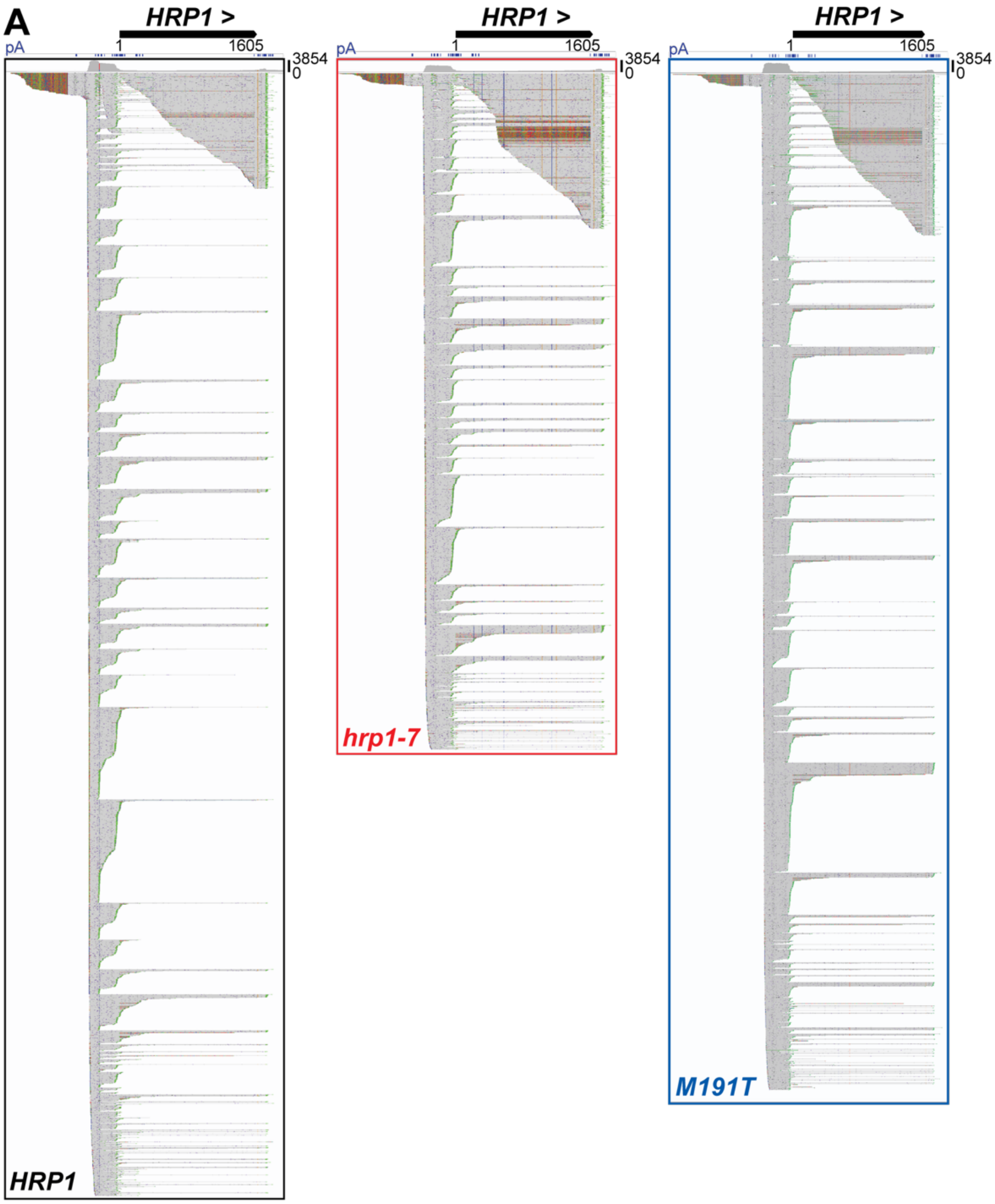


**B**

```
                TSS
-400 TCCTTTCCGCCACTGTAATTAAAAACAAAGGATTGAACAGTTTCGACTAGTTATTTAATTTTGCAGATAGTCAAGTATTAGTACAATATACTGTTATAAA
               TERM1
-300 TTTTTTACGTGTATCTTTTACTTTTTAGTGTTTAATAACTAAAATCTCATACCTTACTAGATTTCTTTCTAACCTTTTTTGCGAAAGAAACAAAAAGAAA
                                                                      TERM2
-200 AAATACGGACAGAAAAGGTATACTCAATAAACAAGAAATTGAAAAAGCGTGCATAATACAAACAATTGGCCCTTTTTTATTCTGTATTAATATTACTGTT

-100 TATATTACAATCTTTTCATAAAAGACGCAAATATTTTATTTATAAAGAACTTTAGCAGTATACACAAGTCTGAGAAAATAGAGAATAAGTTAAATAAGCA

  +1 ATGAGCTCTGACGAAGAAGATTTCAACGACATCTACGGCGATGATAAGCCTACCACTACTGAAGAAGTCAAAAAAGAAGAAGAACAAAATAAGGCTGGCA
```

**Supplemental Figure S3. A)** Unfiltered and unsorted dRNA-seq reads from the indicated strains aligned to the *HRP1* locus. Unaligned bases are shown in color to indicate sequences derived from the pRS313 plasmid (upstream of *HRP1*) and the *KanMX4* cassette (in place of the *HRP1* ORF at the genomic locus). **B)** DNA sequence of the *HRP1* 5'-UTR. The transcription start site region (TSS) and two main attenuation regions (Term 1 and Term 2) are in bold. Numbering is relative to the ATG start codon (+1).

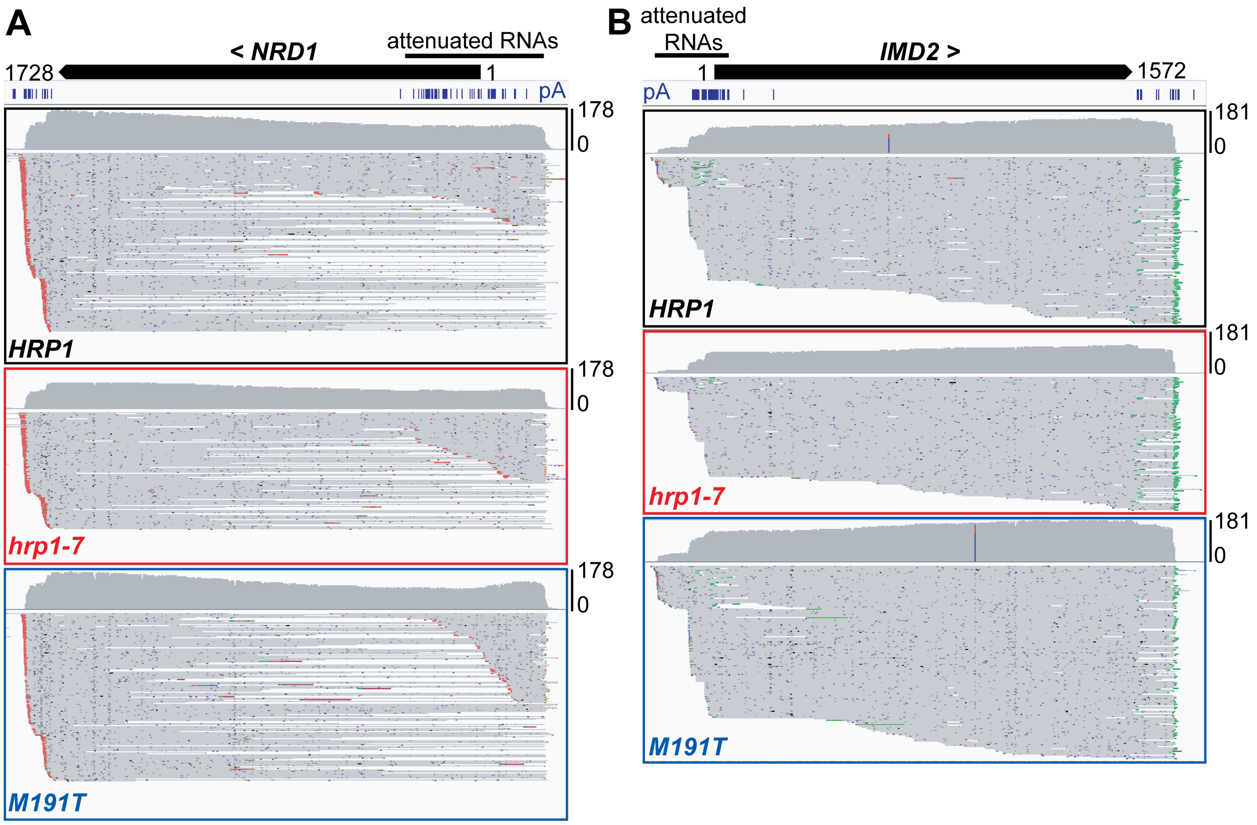


**Supplemental Figure S4. *NRD1* and *IMD2* attenuated transcripts are not as abundant as their mRNAs**. The heavy lines demarcate the ORFs and the lighter lines above demarcate the attenuated transcripts, which are interleaved with the mRNAs. Note that *NRD1* is transcribed leftward and *IMD2* is transcribed rightward. Leftward polyA tails are colored red and rightward are colored green. The *NRD1* attenuated transcripts use the same transcription start site, while the *IMD2* attenuated transcripts use an upstream start site (Kuehner and Brow 2008).

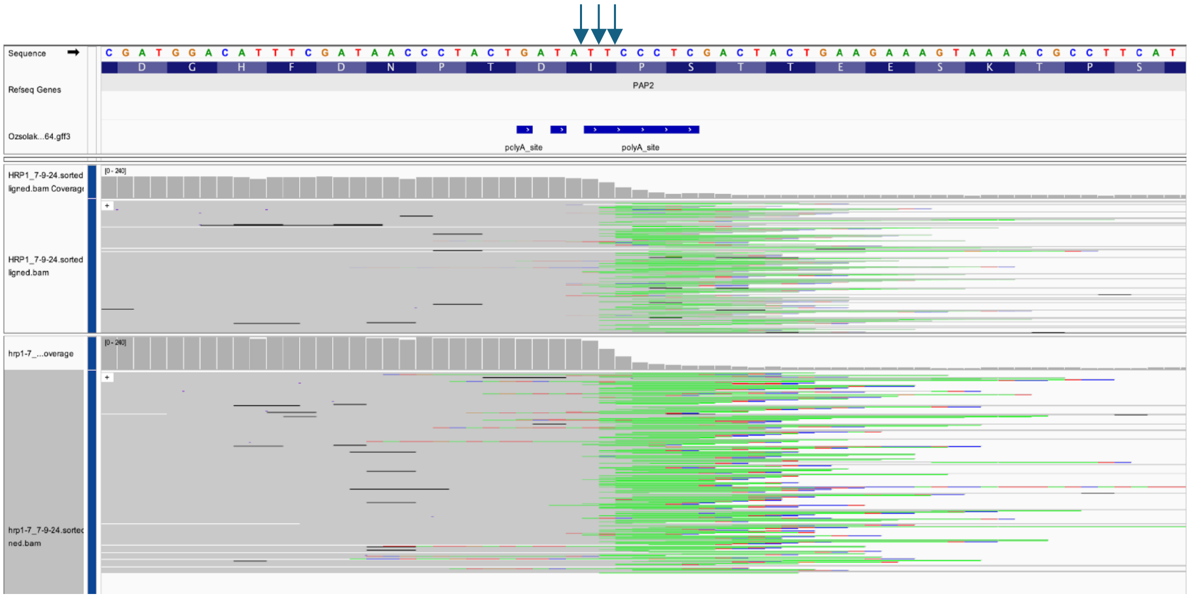


**Supplemental Figure S5.** 3' end positions of truncated *PAP2/TRF4* reads. The most frequent 3' nucleotides before the polyA tail (in green) are indicated with arrows.
